# Recognition of Divergent Viral Substrates by the SARS-CoV-2 Main Protease

**DOI:** 10.1101/2021.04.20.440716

**Authors:** Elizabeth A. MacDonald, Gary Frey, Mark N. Namchuk, Stephen C. Harrison, Stephen M. Hinshaw, Ian W. Windsor

## Abstract

The main protease (M^pro^) of severe acute respiratory syndrome coronavirus 2 (SARS-CoV-2), the cause of coronavirus disease (COVID-19), is an ideal target for pharmaceutical inhibition. It is required for infection, it cleaves the viral polyprotein at multiple sites, and it is conserved among coronaviruses and distinct from human proteases. We present crystal structures of SARS-CoV-2 M^pro^ bound to two viral substrate peptides. The structures show how M^pro^ recognizes substrates and how the peptide sequence can dictate catalytic efficiency by influencing the position of the scissile bond. One peptide, constituting the junction between viral non-structural proteins 8 and 9 (nsp8/9), has P1’ and P2’ residues that are unique among SARS-CoV-2 cleavage sites but conserved among nsp8/9 junctions in coronaviruses. M^pro^ cleaves nsp8/9 inefficiently, and amino acid substitutions at P1’ or P2’ can enhance catalysis. Visualization of M^pro^ with intact substrates provides new templates for antiviral drug design and suggests that the coronavirus lifecycle selects for finely tuned substrate-dependent catalytic parameters.

## TEXT

Developing and stockpiling pan-coronavirus antiviral drugs for pandemic prevention has been a goal since the SARS outbreak of 2003.^1, 2^ The coronavirus main protease (nsp5 or M^pro^) is a conserved drug target and an important focus of these efforts. Hundreds of M^pro^ inhibitors have been reported. Most of these drugs occupy the active site cleft responsible for recognizing the N-terminal half of substrate peptides, and many form covalent bonds to the active site cysteine of M^pro^ (Cys145).^3–7^ A recent crystal structure of the nsp5/6 acyl-enzyme intermediate provides one template for chemical mimicry of this essential catalytic step.^8^ We provide evidence that contacts on both sides of the M^pro^ catalytic site can affect the rate of formation of the covalent complex, a characteristic that could be exploited by new protease inhibitors.

The nsp8/9 junction is a conserved M^pro^ substrate (Figure 1A-B). The nearly invariant Asn residues at P1’ and P2’ are unique within a given coronavirus polyprotein; all other cleavage sites have Gly, Ser, or Ala at P1’, P2’, or both sites.^9^ Cleavage of nsp8/9 is slow but required for replication of the closely-related Murine Hepatitis Virus.^10^ Indeed, a recently-determined cryo-EM structure shows that the N-terminus of nsp9 contacts nsp12, a core component of the viral RNA polymerase.^11^ In this context, the nsp8/9 P1’ to P3’ residues contribute to a binding site for a nucleotide that is transferred to the amino terminus of the P1’ residue.^12^ Therefore, the nsp8/9 junction has evolved to satisfy two evolutionary constraints required for viral replication: it must be cleaved in the M^pro^ active site, and it must serve as a substrate in a nucleotide monophosphate transfer reaction catalyzed by the nsp12 protein. We have used X-ray crystallography to study nsp8/9 and nsp4/5 recognition by The structures show unique features of the M^pro^·nsp8/9 complex constrained by natural selection and highlight the importance of P1’-P3’ residues in catalysis.

**Figure 1.**
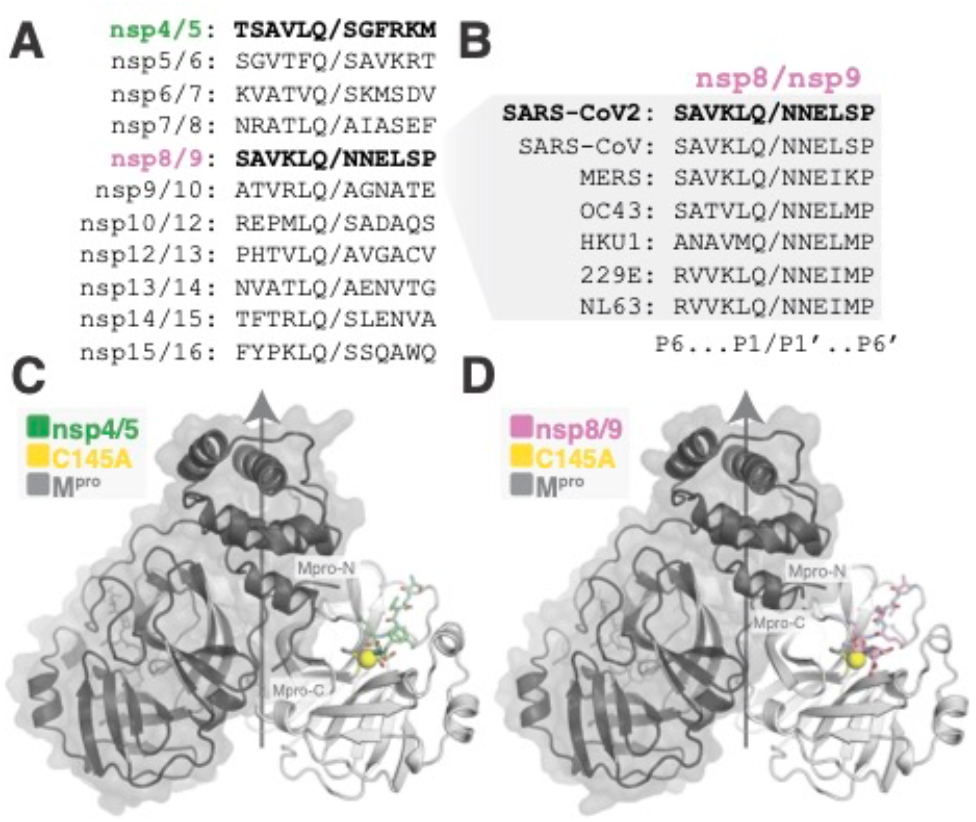
Viral substrates and structures of substrate-bound M^pro^. (A) Protein sequence alignment of the eleven SARS-CoV-2 M^pro^ cleavage sites required for maturation of SARS-CoV2. (B) Protein sequence alignment of nsp8/9 M^pro^ cleavage sites from representative coronaviruses. (C-D) Crystal structures showing nsp4/5 (C) or nsp8/9 (D) substrate engagement by M^pro^. Protease protomers are colored dark or light gray. A single peptide is colored for each, though peptide occupies all active sites in both determined structures. The vertical arrow marks the two-fold axis of the M^pro^ dimer.

To study M^pro^ activity, we monitored cleavage of labeled substrate peptides *in vitro* and derived Michaelis-Menten parameters describing the reactions. M^pro^ cleavage of nsp8/9 is less efficient than cleavage of nsp4/5 (36-fold decrease in *k*_cat_/*K*_M_; Table 1).^13^ We also sought to understand the influence of the Asn residues at the P1’ and P2’ sites of the nsp8/9 substrate (Table 1). Alanine substitutions at either position approximately doubled the catalytic efficiency. The P1’ Asn-to-Ala substitution lowered *K*_M_ and raised *k*_cat_, while the P2’ substitution only raised

**Table 1.** Catalytic efficiencies for M^pro^ substrates and analogs

| Substrate | Sequence<sup>a</sup> | $k_{\text{cat}}$ (s<sup>-1</sup>) | $K_M$ (μM) | $k_{\text{cat}}/K_M$ (M<sup>-1</sup>s<sup>-1</sup>) | Fold change<sup>b</sup> |
| --- | --- | --- | --- | --- | --- |
| nsp4/5 | TSAVLQ/SGFRKM | $0.52 \pm 0.07$ | $41 \pm 9$ | $1.3 \pm 0.3 \times 10^4$ | — |
| nsp8/9 | RVVKLQ/NNELMP | $0.013 \pm 0.001$ | $36 \pm 6$ | $3.6 \pm 0.7 \times 10^2$ | 1.0 |
| nsp8/9 N1'A | RVVKLQ/aNELMP | $0.022 \pm 0.001$ | $22 \pm 3$ | $1.0 \pm 0.1 \times 10^3$ | 2.9 |
| nsp8/9 N2'A | RVVKLQ/NaELMP | $0.034 \pm 0.002$ | $46 \pm 5$ | $7.5 \pm 0.8 \times 10^2$ | 2.1 |
| nsp8/9 N1'D | RVVKLQ/dNELMP | — | — | — | — |
| nsp8/9 N2'D | RVVKLQ/NdELMP | $0.0029 \pm 0.0001$ | $19 \pm 1$ | $1.6 \pm 1.2 \times 10^2$ | 0.4 |

<sup>a</sup>Lys-DABCYL and Glu-EDANS and are appended to the N- and C-termini. Residues that differ from the wild-type sequence are lower case. <sup>b</sup>Fold change = $(k_{\text{cat}}/K_M)_{\text{nsp8/9 analog}} / (k_{\text{cat}}/K_M)_{\text{nsp8/9}}$

*k*_cat_. Installation of an isosteric Asp residue at the P1’ position completely abrogated activity, while the analogous Asn-to-Asp substitution at P2’ diminished but did not abrogate activity. We suspect that placing additional negative charge near the active site raises the energetic barrier to attaining the oxyanion transition states.^14, 15^

Differences in *k*_cat_ for the peptide substrates tested dominated the small changes in *K*_M_ and drove the observed changes in *k*_cat_/*K*_M_. The P5-P1 residues were constant for nsp8/9 and its derivatives, ruling out acyl-enzyme hydrolysis as the step that determines *k*_cat_. Therefore, either formation of the enzyme-substrate complex or conversion to the acyl-enzyme intermediate must limit *k*_cat_ for the nsp8/9 substrate, and similar *K*_M_ values imply the latter is true. Unlabeled nsp8/9 did not inhibit cleavage of labeled nsp4/5 at concentrations up to 250 μM, while the control inhibitor MG-132 produced a *K*_i_ of 740 nM (Figure S2).^5^ Its inability to interfere with nsp4/5 cleavage confirms that nsp8/9 is a poor M^pro^ substrate relative to nsp4/5. Similar competition among the 11 viral M^pro^ substrates occurs during virus replication.

To understand the structural basis for differential cleavage efficiency, we determined crystal structures of M^pro^ bound to the nsp4/5 and nsp8/9 substrates (Figure 1C-D). The active site Cys145Ala mutation trapped the intact substrates and enabled visualization of the P’ residues (Figure S3). A principal difference between the nsp4/5 and nsp8/9 substrates is the position of the scissile bond. The nsp8/9 P1 Cα is translated 0.4 Å along the C=O axis relative to nsp4/5 (Figure 2). Nevertheless, 11 conserved hydrogen bonds occur between M^pro^ and each of the substrates (Figure 3, white dotted lines). Eight contacts between the peptide backbones of enzyme and substrate are shared among SARS-CoV nsp4/5, PEDV nsp4/5,^16, 17^ and the two SARS-CoV-2 peptides reported here. M^pro^ Gly143 and Ala145 mainchain amides form the oxyanion hole by donating a pair of hydrogen bonds to the scissile P1 carbonyl oxygen, which stabilizes the developing negative charge during covalent catalysis. His163 and the mainchain carbonyl of Phe140 make hydrogen bonds with the invariant sidechain of the P1 Gln, and Asn142 contacts the P1 Gln through a conserved water bridge. Neither SARS-CoV nor PEDV M^pro^·nsp4/5 complexes show the hydrogen bonds observed with Asn142 in the SARS-CoV2 substrate complexes.^16, 17^

**Figure 2.**
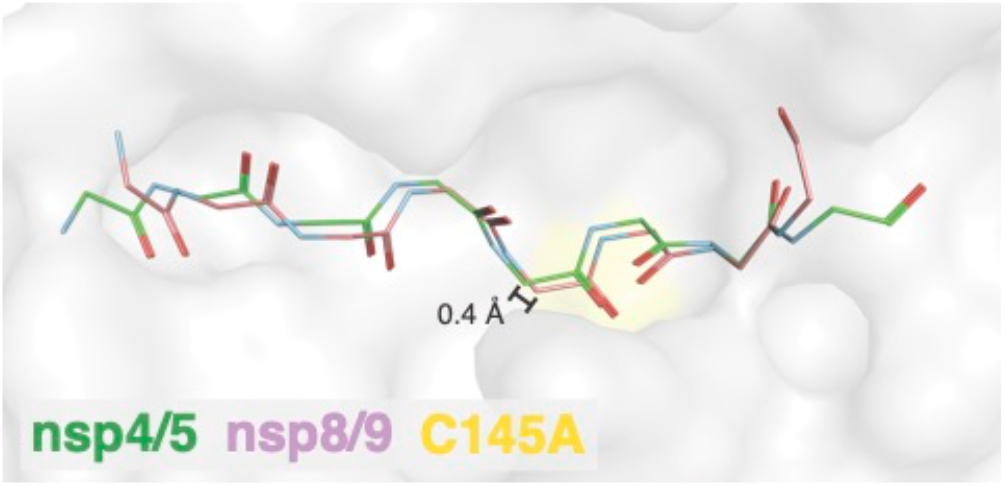
Backbone view showing shifted M^pro^ substrates. The catalytic site mutation M^pro^ Cys145Ala is yellow (surface). Distance between P1 Cα for nsp4/5 (green) and nsp8/9 (pink) is given.

**Figure 3.**
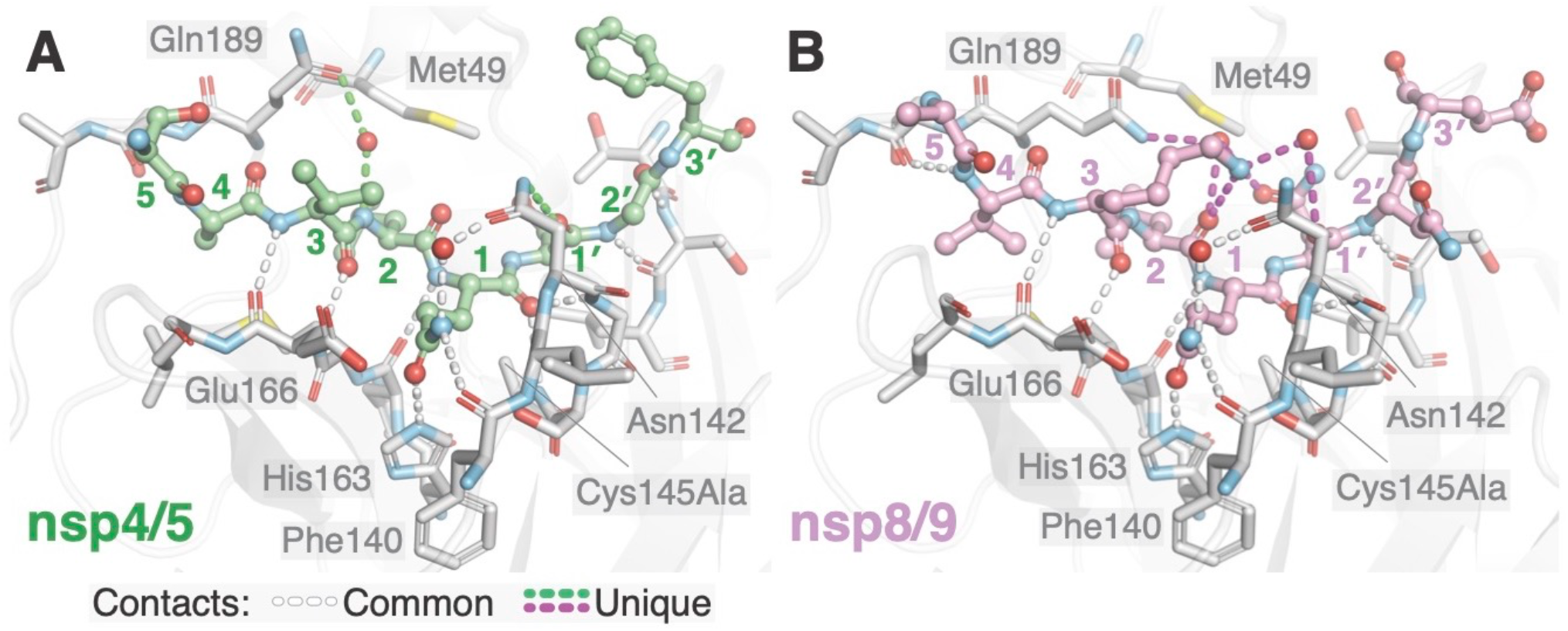
Differential recognition of nsp4/5 and nsp8/9 substrates by M^pro^. Identical views of nsp4/5 (A) and nsp8/9 (B) substrates in the M^pro^ Cys145Ala active site. Substrate peptide P and P’ residues are labeled with colored numbers. Key M^pro^ residues mentioned in the text are labeled. Conserved hydrogen bonds enabling M^pro^ recognition of substrate mainchain and P1 Gln side chain atoms are shown as white dotted lines. The hydrogen bonds between M^pro^ and nsp4/5 (green) differ from those with nsp8/9 (magenta). M^pro^ Asn142 and Gln189 contact the nsp4/5 substrate through bound water molecules, while the same M^pro^ residues contact the nsp8/9 peptide at distinct sites through networked water molecules.

Substrate interactions with the M^pro^ Asn142 and Gln189 sidechains distinguish nsp4/5 and nsp8/9 recognition (Figure 3, green and magenta dotted lines). M^pro^ Asn142 forms a hydrogen bond with the nsp4/5 P1’ backbone carbonyl oxygen, and M^pro^ Gln189 forms a water bridge with the nsp4/5 P2 amide nitrogen. In contrast, M^pro^ Gln189 engages the nsp8/9 P3 and P1’ side chains via an ordered water molecule. In addition to these contacts, the ordered waters found in the nsp8/9-bound structure could donate hydrogen bonds to the P1’ and P2 mainchain carbonyl oxygens. Finally, the nsp8/9 P3 Lys forms a hydrogen bond with the P2 carbonyl. This interaction and the ordered waters described above likely account for subtle bending of the nsp8/9 substrate P-fragment away from the active site cleft relative to the nsp4/5 peptide.

The peptide recognition described above produces the different catalytic efficiencies associated with cleavage of the nsp4/5 and nsp8/9 substrates. The near-invariant nsp8/9 P1’ and P2’ Asn side chains are bulkier than the P1’ (Ser/Ala) and P2’ (Gly/Ala) side chains of other M^pro^ substrates, though there is greater tolerance for P2’ diversity.^18^ The nsp8/9 P1’ Asn reaches more deeply into the S’ subsite than the nsp4/5 P1’ Ser and therefore likely restrains the P’ peptide to a greater degree. M^pro^ Asn142 and Gly143 coordinate the nsp8/9 P2’ residue through peptide backbone interactions, and similar interactions position the nsp4/5 P2’ Gly (Figure 4A-B). Overall, the bulkier nsp8/9 Asn side chains in the S1’ and S2’ subsites shift nsp8/9 relative to nsp4/5 (Figure 2). The resulting alignment with the M^pro^ cysteine nucleophile differs from that of nsp4/5 and provides an explanation for reduced catalytic efficiency that is partially restored by alanine substitution. That the restoration is incomplete relative to nsp4/5 suggests that both P- and P’-substrate fragments align the scissile bond.

**Figure 4.**
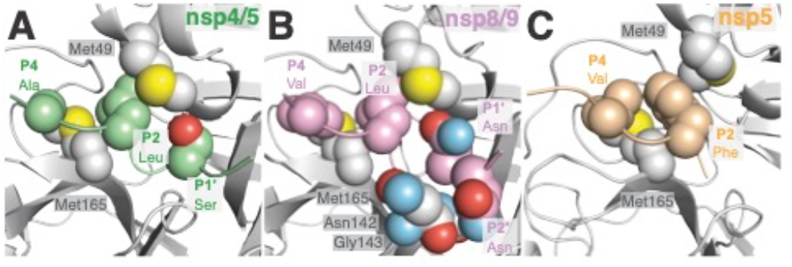
Steric effects that influence M^pro^ recognition and activity. Key atoms dictating the shape complementarity between M^pro^ subsites and corresponding side chains from: (A) nsp4/5, (B) nsp8/9, and (C) nsp5 (acyl-enzyme intermediate; PDB 7KHP).

Hydrophobic interactions dictate recognition of N-terminal substrate fragments (P residues, excluding the invariant P1 Gln). M^pro^ Met49 and Met165 define the S4, S2, and S1’ subsites (Figure 4). The nsp4/5 P3 Ala is smaller than the ns8/9 P3 Val, allowing the P4 Ala of nsp4/5 to sit more deeply in the S4 subsite. This produces a 1.0 Å shift between the nsp4/5 and nsp8/9 P3 residues (measured from corresponding Cα atoms). A recent crystal structure shows that the intact nsp5/6 substrate is also shifted relative to nsp4/5 due to a bulky Phe at the P2 position (Figure 4C).^8^ Indeed, M^pro^ cleavage is most efficient for peptides bearing P2 Leu and less efficient for those bearing P2 Phe.^19^ Like nsp8/9, cleavage of SARS nsp5/6 depends more heavily on P’ recognition than does nsp4/5.^20, 21^ The nsp8/9 P3 Lys might also limit catalysis. Water bridges connect its terminal nitrogen (Nζ) with the nsp8/9 P1’ Asn (mentioned above), and the resulting conformation could slow peptide accommodation to the M^pro^ active site. Therefore, the diverse M^pro^-substrate interactions can fine-tune substrate geometry and position the scissile bond to adjust substrate-specific activity.^22^

The structures we have resolved show how M^pro^ active site plasticity and substrate evolution can tune catalysis. Slow cleavage of the nsp8/9 junction, which is observed among disparate coronaviruses, might be a selected trait required for coordinated assembly of the RNA replication machinery.^9, 13, 23, 24^ Inability of the nsp8/9 substrate to inhibit nsp4/5 cleavage is likely important for maturation of the viral polyprotein. The need for the nsp8/9 junction to support both M^pro^ cleavage and nsp12 binding (and subsequent nucleotide monophosphate acceptance) accounts for the near-invariance of the P1-P2’ residues. The sequence is therefore a compromise that satisfies the requirements of two unrelated catalytic mechanisms, and mimicry of the nsp8/9 junction presents a unique opportunity to chemically inhibit both M^pro^ and the viral polymerase.

The structures also present templates for new protease inhibitor scaffolds. In particular, that the nsp8/9 P3 sidechain can fold back to contact P1’ suggests macrocyclic inhibitors could mimic this interaction. Similar strategies have been pursued for Hepatitis C NS3, HIV-1, and Rhinovirus 3C proteases.^25–27^ Kinetic analyses of nsp8/9 and its variants suggests that inhibitor P1’ and P2’ site contacts could influence formation of covalent inhibitor-enzyme adducts. α-ketoamide warheads, which have not been exhaustively explored as ligands for M^pro^ Cys145, are good candidates for this objective.

## ASSOCIATED CONTENT

### Supporting Information

Supporting Information document contains methods, inhibition experiments, and crystallographic information. Crystallographic data has been deposited at the PDB with associated codes 7MGR (M^pro^·nsp8/9) and 7MGS (M^pro^·nsp4/5).

## AUTHOR INFORMATION

### Author Contributions

The manuscript was written through contributions of all authors. All authors have given approval to the final version of the manuscript.

## ACKNOWLEDGMENTS

Enzyme assays were performed at the Institute for Chemistry and Chemical Biology (ICCB-Longwood) at Harvard Medical School. We thank Linlin Zhang (Hilgenfeld Laboratory, University of Lübeck) for providing the codon-optimized Mpro expression plasmid. X-ray diffraction data were collected at the Advanced Photon Source (APS) on NE-CAT beamline 24-IDC.

