## Supplemental Information for "Recognition of Divergent Viral Substrates by the SARS-CoV-2 Main Protease"

| Contents | Page |
|---|---|
| Table of Contents | S1 |
| Materials and Methods | S2 |
| References | S4 |
| Figure S1. Michaelis-Menten parameters from M$^{pro}$ activity assays | S5 |
| Figure S2. Evaluation of inhibition of M$^{pro}$ by MG-132 and nsp8/9 substrate | S6 |
| Figure S3. Fo-Fc omit map density for substrates | S7 |
| Table S1. Crystallographic data collection and model statistics | S8 |

### MATERIALS AND METHODS

#### Protein Expression and Purification

A plasmid containing codon-optimized SARS-CoV-2 main protease ($M^{pro}$) was a gift from Zhang *et al.*[1] $M^{pro}$ expressed and purified from *E. coli* carrying this plasmid was used for peptide cleavage assays. For production of inactive $M^{pro}$, an insert coding for SUMO-$M^{pro}$ was prepared by overlap extension PCR and inserted into pLIC 2B-T (Addgene #29666) by ligation-independent cloning. The codon-optimized $M^{pro}$ gene was then mutagenized by overlap extension PCR to create $M^{pro}$ Cys145Ala (fwd - ggcagcttccttaatggcagcGCGGGTTCGGTGGGCTTTAACATCG; rev - CGATGTTAAAGCCCACCGAACCCGCgctgccattaaggaagctgcc) before reinsertion into the parent vector. The resulting construct codes for a His6-TEV site-SUMO-$M^{pro}$-PreScission site-His6 fusion protein. Ulp1 cleavage produces the native $M^{pro}$ N-terminus. PreScission protease cleavage produces the native $M^{pro}$ C-terminus.[2]

Wild type or $M^{pro}$ Cys145Ala plasmids were transformed into Rosetta 2(DE3)pLysS *E. coli* (Novagen) and grown overnight. 8mL of the overnight culture was used to seed flasks of 1 L 2XYT (~12 L total culture volume). The cells were grown at 37 °C shaking at 225 RPM until they reached an optical density at 600 nm of 1.0. Protein expression was then induced with 400 μM isopropyl β-d-1-thiogalactopyranoside (IPTG), and the temperature was changed to 18 °C. After ~18 hours, the cells were pelleted at 4°C for 30 minutes and resuspended in 5 mL per liter of culture in lysis buffer (20 mM Tris pH 8.0, 150 mM NaCl, 10 mM Imidazole, and 1 mM β-mercaptoethanol (BME)). Cell pellets were frozen and stored at -80 °C until needed.

Cells were lysed by sonication, and then the lysate was centrifuged at 19,000 RPM for 1 hour in a Beckman JA-20 rotor. Typically, the soluble fraction 12 L of culture was incubated with 5 mL TALON cobalt resin (Takara) at 4 °C for one hour. Cobalt resin was washed with 50 mL Lysis Buffer in 10 mL increments, then with 10 mL Wash Buffer (20 mM Tris, pH 8; 50 mM NaCl; 10 mM imidazole; 1 mM BME). SUMO-$M^{pro}$ Cys145Ala was eluted from the cobalt resin in 50 mL Elution Buffer in 10 mL increments (20 mM Tris, pH 8; 50 mM NaCl; 400 mM Imidazole; 1 mM BME). The SUMO tag was removed by incubating the pooled eluate with Ulp1 overnight at 4 °C while dialyzing against 20 mM Tris, pH 8; 50 mM NaCl; and 1 mM BME. The cleaved protein was passed over cobalt resin to remove uncleaved fusion protein, Ulp1, and SUMO. Additional contaminants and remaining SUMO were removed using sequential cation and anion exchange chromatography by linking the columns directly (HiTrap Q HP and HiTrap SP HP, GE). Fully cleaved $M^{pro}$ Cys145Ala did not bind either column and was collected in the flowthrough. $M^{pro}$ Cys145Ala was concentrated to 20 mg/mL by ultrafiltration. The buffer was gradually exchanged to crystallization buffer during this step (20 mM Tris pH 8; 150 mM NaCl; and 1 mM BME).

#### Crystallization and Structure Determination

Substrate peptides corresponding to P4-P5′ of the nsp4/5 (AVLQSGFRK) and nsp8/9 (AVKLQNNEL) cleavage sequences were synthesized by Biomatik Peptide Synthesis Services using solid phase peptide synthesis. Peptides were resuspended to a final concentration of 6 mM in sterile deionized water. Co-crystals of $M^{pro}$ Cys145Ala and substrate peptides were obtained by hanging-drop vapor diffusion. Equal volumes of 6 mM peptide and 20 mg/mL $M^{pro}$ Cys145Ala were mixed to produce 10 mg/mL $M^{pro}$ Cys145Ala with a 10-fold molar excess of substrate. Mixtures were incubated on ice for one hour to allow substrate binding to the enzyme. The $M^{pro}$ complexes were mixed 1:1 (v/v) with mother liquor. $M^{pro}$ Cys145Ala·nsp4/5 crystals were grown in 9% Polyethylene Glycol (PEG) 6000 and 0.1 M MES, pH 6.5 at 18 °C. Rod-shaped crystals formed after 24 hours. $M^{pro}$ Cys145Ala·nsp8/9 crystals were grown in 15% PEG 4000; 0.1 M Tris, pH 8; and 5% Dimethyl Sulfoxide (DMSO), at 4 °C. Clusters of cube-shaped crystals formed after 7 days. Crystals were transferred to a cryoprotectant containing a modified mother liquor containing 1.2-fold concentrated components, 1.5 mM substrate peptide, and 20% PEG 400 before flash-freezing in liquid nitrogen.

Diffraction experiments were conducted at the Advanced Photon Source (APS) of Argonne National Lab on NE-CAT beamline 24ID-C. Datasets were indexed, integrated, and scaled using Xray Detector Software (XDS).[3] Phases were determined by molecular replacement with Phaser as implemented in Phenix using PDB entry 6Y2E as a search model.[1, 4, 5] Models were refined using Phenix Refine. Peptide models were built using Coot and validated with MolProbity.[5-7]

**Enzyme Kinetics**

Förster resonance energy transfer (FRET) substrate peptides labeled with N-terminal fluorophore Dabcyl and C-terminal quencher Edans were purchased from New England Peptide. FRET substrate peptides represent the wildtype nsp4/5 cleavage sequence (nsp4/5 WT, Dabcyl-KTSAVLQSGFRKME-Edans), nsp8/9 cleavage sequence (nsp8/9 WT, Dabcyl-KRVVKLQNNEIME-Edans), nsp8/9 cleavage sequence with P1' Alanine (1A, Dabcyl-KRVVKLQ**A**NEIME-Edans), nsp8/9 cleavage sequence with P2′ Alanine (2A, Dabcyl-KRVVKLQN**A**EIME-Edans), nsp8/9 cleavage sequence with P1′ Aspartate (2A, Dabcyl-KRVVKLQ**D**NEIME-Edans), and nsp8/9 cleavage sequence with P2′ Aspartate (2A, Dabcyl-KRVVKLQN**D**EIME-Edans). The peptides were dissolved at 10 mM in DMSO.

All cleavage assays were performed at 25 °C in 20 mM Tris(HCl) pH 8.0, 100 mM NaCl, 0.5 mM EDTA, 1 mM TCEP. For kinetic analysis, 10 µl of assay buffer was dispensed into black 384 well microtiter plates (Corning 3820) using a Thermo Scientific Multidrop Combi. Varying concentrations of substrate were dispensed using an HP D300 Digital Dispenser. The final concentration of DMSO in the assay was 0.5%. The reaction was initiated with the addition of 10 µl of $M^{pro}$ and fluorescence monitored on a Flexstation 3 plate reader (Molecular Devices) with excitation at 360 nm and emission at 460 nm. For results reported in Table 1, [$M^{pro}$] = 0.25 µM for nsp4/5 experiments, and [$M^{pro}$] = 0.4 µM for nsp8/9 experiments.

Relative fluorescence units (RFU) were calibrated to product concentration by measuring the fluorescence of a range of concentrations of product. Linear regression with Prism 6 (Graphpad Software) determined a slope of 14.4 RFU/µM, which was used to convert initial velocities form RFU/s to nM/s. Initial velocities were calculated by linear regression with Prism 6. Mean initial velocities and standard deviations from triplicate measurements versus substrate concentration were fit with the "kcat" built-in function by non-linear regression analysis using Prism 6. This version of the Michael-Mention equation requires an input of enzyme concentration as a fixed parameter and fits the turnover number ($k_{cat}$) and the Michealis constant ($K_M$). Values from the non-linear regression fitted parameters are listed in the main text along with the standard error. Initial velocities were obtained with the same approach as the Michealis-Menten kinetics and values were fit with "one site – total" function by non-linear regression in Prism 6.

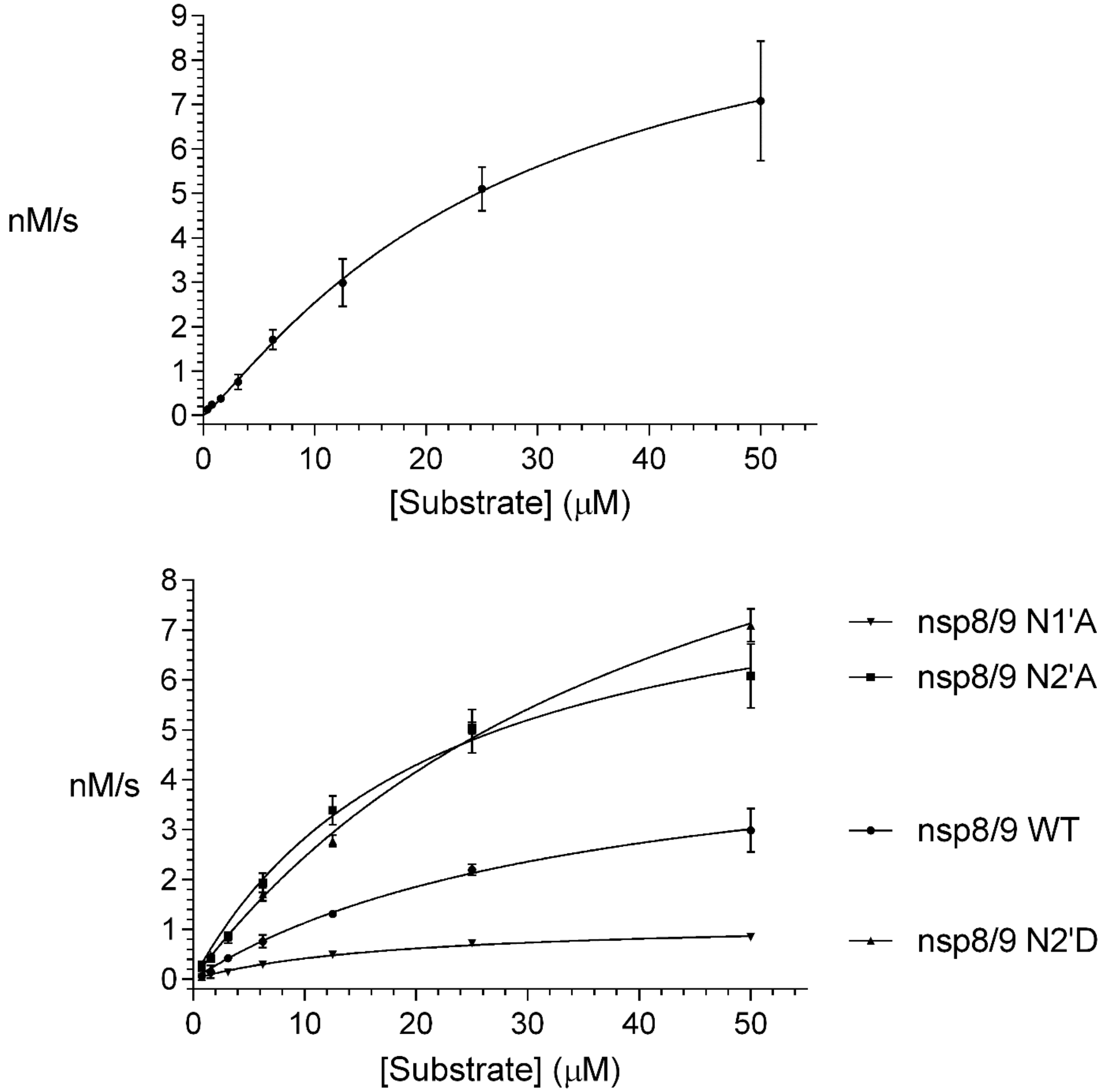


**Figure S1.** Michaelis-Menten parameters from M$^{pro}$ activity assays. Substrate concentration dependence of initial velocities for nsp4/5 cleavage (top) and for nsp8/9 cleavage (including associated analog; bottom) are shown.

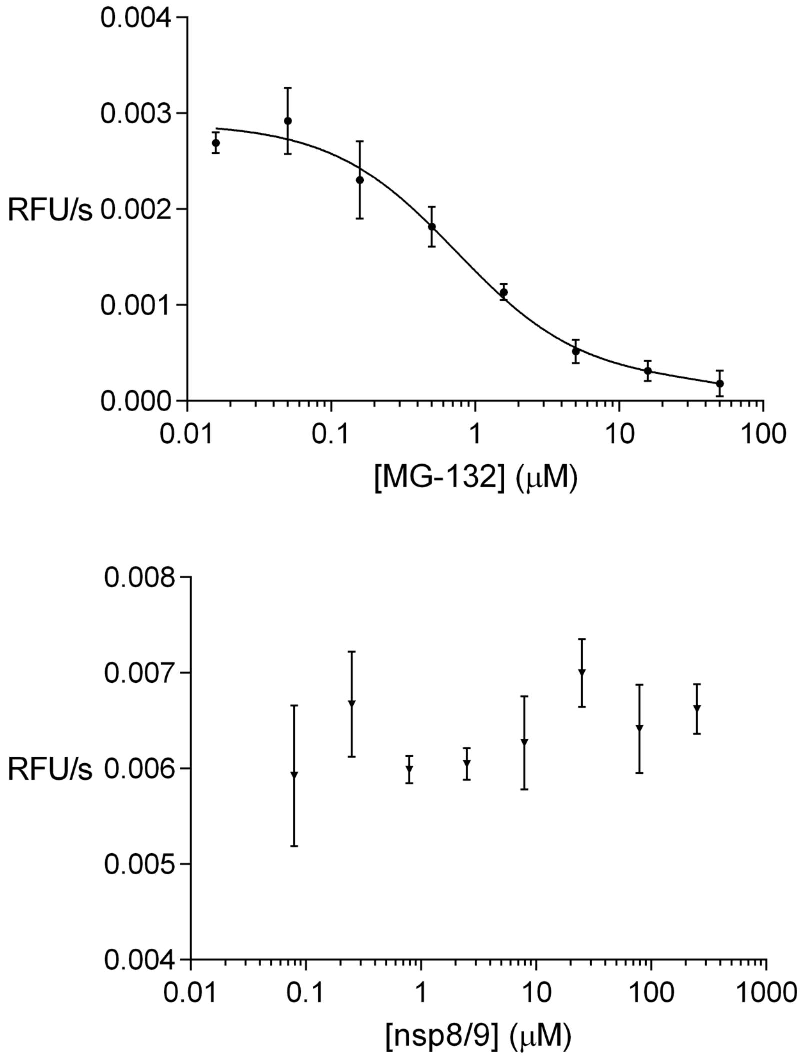


**Figure S2.** Evaluation of inhibition of $M^{pro}$ by MG-132 and nsp8/9 substrate. The activity of $M^{pro}$ towards the EDANS/DABCYL-labeled nsp4/5 substrate was measured in the presence of varied concentrations of the covalent inhibitor MG-132 (top) or unlabeled nsp8/9 substrate (bottom). The value of $K_i$ determined for MG-132 was 740 nM. No dosage dependent reduction in activity was observed upon addition of nsp8/9 substrate. 10 nM $M^{pro}$ was used for the MG-132 experiments, and 20 nM $M^{pro}$ was used for the nsp8/9 experiments.

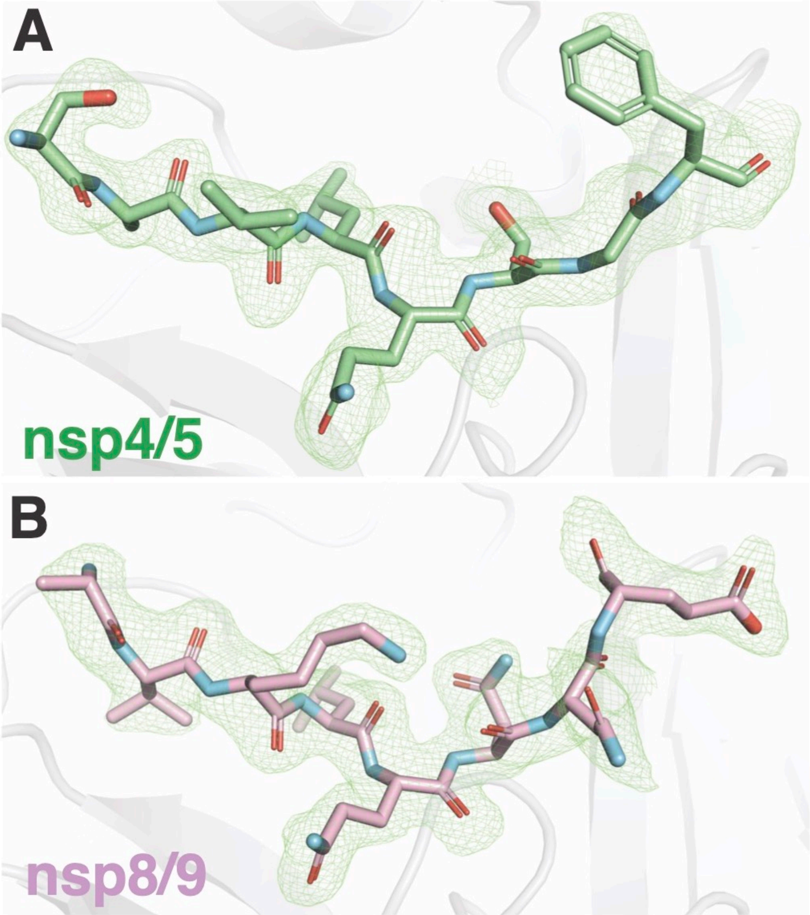


**Figure S3.** Fo-Fc omit map density for substrates. Density is contoured at 1.5σ for nsp4/5 (A) and 2.5σ nsp8/9 (B). Peptides are colored as indicated, density is carved within 2.0 Å of the peptides, and $M^{pro}$ is gray.

| | SARS-CoV-2 Mpro + nsp4/5 PDB 7MGS | SARS-CoV-2 Mpro + nsp8/9 PDB 7MGR |
|---|---|---|
| **Wavelength** | 0.979180 | 0.979110 |
| **Resolution range** | 104.6 - 1.84 (1.906 - 1.84) | 45.1 - 1.94 (2.009 - 1.94) |
| **Space group** | P 21 21 2 | C 1 2 1 |
| **Unit cell** | 45.771 63.312 104.646 90 90 90 | 99.398 81.982 52.123 90 114.837 90 |
| **Total reflections** | 353281 (36649) | 188911 (19439) |
| **Unique reflections** | 27123 (2501) | 27966 (2608) |
| **Multiplicity** | 13.0 (13.7) | 6.8 (6.9) |
| **Completeness (%)** | 91.53 (93.46) | 98.54 (89.94) |
| **Mean I/sigma(I)** | 18.00 (3.96) | 11.47 (2.54) |
| **Wilson B-factor** | 36.57 | 38.64 |
| **R-merge** | 0.09926 (0.94) | 0.08605 (0.4647) |
| **R-meas** | 0.1035 (0.9763) | 0.0933 (0.5014) |
| **R-pim** | 0.02893 (0.2612) | 0.03565 (0.1868) |
| **CC1/2** | 0.998 (0.942) | 0.997 (0.965) |
| **CC*** | 1 (0.985) | 0.999 (0.991) |
| **Reflections for refinement** | 24844 (2499) | 27728 (2539) |
| **Reflections used for R-free** | 1996 (199) | 1988 (181) |
| **R-work** | 0.1934 (0.3113) | 0.1939 (0.2976) |
| **R-free** | 0.2272 (0.3571) | 0.2351 (0.3635) |
| **CC(work)** | 0.962 (0.842) | 0.963 (0.895) |
| **CC(free)** | 0.955 (0.789) | 0.953 (0.855) |
| **Number non-hydrogen atoms** | 2497 | 2517 |
| **macromolecules** | 2413 | 2403 |
| **ligands** | 1 | |
| **solvent** | 83 | 114 |
| **Protein residues** | 313 | 309 |
| **RMS(bonds)** | 0.009 | 0.011 |
| **RMS(angles)** | 1.19 | 2.05 |
| **Ramachandran favored (%)** | 97.41 | 97.38 |
| **Ramachandran allowed (%)** | 2.27 | 2.62 |
| **Ramachandran outliers (%)** | 0.32 | 0 |
| **Rotamer outliers (%)** | 0.75 | 0.75 |
| **Clashscore** | 2.93 | 6.51 |
| **Average B-factor** | 39.73 | 48.39 |
| **macromolecules** | 39.63 | 48.36 |
| **ligands** | 56.41 | |
| **solvent** | 42.23 | 49.08 |

Table S1. Crystallographic data collection and model statistics
